# mobsim: An R package for the simulation and measurement of biodiversity across spatial scales

**DOI:** 10.1101/209502

**Authors:** Felix May, Katharina Gerstner, Dan McGlinn, Xiao Xiao, Jonathan M. Chase

**Affiliations:** German Centre for Integrative Biodiversity Research (iDiv) Halle-Jena-Leipzig, Deutscher Platz 5e, D-04103 Leipzig, Germany; Institute of Computer Science, Martin-Luther University Halle-Wittenberg, D-06099 Halle (Saale), Germany; Biology Department, College of Charleston, Charleston, SC, USA; School of Biology and Ecology, and Senator George J. Mitchell Center for Sustainability Solutions, University of Maine, Orono, ME, USA 04469

**Keywords:** diversity indices, sampling, scale-dependence, simulation, species-abundance distribution, species-accumulation curve, species-area relationship, rarefaction curve

## Abstract

1. Estimating biodiversity and its changes in space and time poses serious methodological challenges. First, there has been a long debate on how to quantify biodiversity, and second, measurements of biodiversity change are scale-dependent. Therefore comparisons of biodiversity metrics between communities are ideally carried out across scales. Simulation can be used to study the utility of biodiversity metrics across scales, but most approaches are system specific and plagued by large parameter spaces and therefore cumbersome to use and interpret. However, realistic spatial biodiversity patterns can be generated without reference to ecological processes, which suggests a simple simulation framework could provide an important tool for ecologists.

2. Here, we present the R package `mobsim` that allows users to simulate the abundances and the spatial distribution of individuals of different species. Users can define key properties of communities, including the total numbers of individuals and species, the relative abundance distribution, and the degree of spatial aggregation. Furthermore, the package provides functions that derive biodiversity patterns from simulated communities, or from observed data, as well as functions that simulate different sampling designs.

3. We show several example applications of the package. First, we illustrate how species rarefaction and accumulation curves can be used to disentangle changes in the fundamental biodiversity components: (i) total abundance, (ii) relative abundance distribution, (iii) and species aggregation. Second, we demonstrate how mobsim can be used to assess the performance of species-richness estimators. The latter indicates how spatial aggregation challenges classical non-spatial species-richness estimators.

4. `mobsim` allows the simulation and analysis of a large range of biodiversity scenarios and sampling designs in an efficient and comprehensive way. The simplicity and control provided by the package can also make it a useful didactic tool. The combination of controlled simulations and their analysis will facilitate a more rigorous interpretation of real world data that exhibit sampling effects and scale-dependence.

## Introduction

Understanding how biodiversity varies in space and time poses one of the greatest challenges in ecology. One reason this challenge is difficult to overcome is that observed biodiversity changes depend on spatial scale (Rosenzweig 1995; Rahbek 2005) and on the specific biodiversity measure used (reviewed in Magurran & McGill 2011). Reasons for the complexity in measuring biodiversity across scales are twofold. First, any measure of biodiversity (e.g., species richness, Shannon, or Simpson diversity) transforms the numbers of individuals and species in a given sample into a single univariate metric that necessarily only captures a portion of information about the underlying abundances and spatial distribution. Second, biodiversity measures vary non-linearly with spatial scale and thus any comparisons among two or more sites will typically be highly scale-dependent (i.e., their difference or ratio will depend on the scale in which it is measured) (Chase & Knight 2013). Despite continued discussion approaches for estimating and comparing diversity measures (e. g. Jost 2006; Colwell *et al*. 2012), no single measure can capture all of the relevant information and multiple measures and approaches will provide more complete information about biodiversity and its change. All biodiversity patterns, including local and regional measures of diversity (α, γ-diversities) and their scaling relationships (measures of β-diversity) depend on three biodiversity components, namely (i) the total abundance of individuals, (ii) relative species abundance distribution, and (iii) the spatial distributions of individuals and species (He & Legendre 2002; McGill 2011). Although we focus on taxonomic diversity measures here, the same issues apply for measurements of functional and (phylo-)genetic diversity (Chao, Chiu & Jost 2014).

Here, we introduce the software package `mobsim` to facilitate understanding and interpretation of biodiversity changes across scales. `mobsim` includes spatially explicit simulation tools, which allow user-defined manipulations of the three biodiversity components. Of course, in nature the components emerge from species traits and dynamic ecological processes such as competition, dispersal limitation, or habitat filtering. However, we suggest that direct simulations and manipulations of the emerging biodiversity components provide generality that embraces different taxa and ecosystems.

Often, when discussing processes of sampling and resulting biodiversity measurements, analogies such as pulling jellybeans from a jar are used (e.g., Gotelli & Colwell 2001). These analogies also provide useful pedagogical tools for teaching biodiversity concepts (e.g., Heard 2016). mobsim simulates individuals of different species (i.e., the proverbial “jelly beans”) in a spatially explicit landscape and thus allows studying the influence of sampling and scale, as well as the interrelatedness between different biodiversity descriptors and patterns in a comprehensive and efficient way. The package provides functions for three purposes (1) the simulation of communities in space, (2) the analysis of biodiversity patterns, and (3) the simulation and assessment of different sampling designs (Fig. 1, Table 1).

Specifically, in spatially-explicit simulations users define the total number of individuals, the species richness, the shape and evenness of the relative species-abundance distribution, and the intraspecific aggregation of species. Functions for the analysis of biodiversity patterns, such as rarefaction-curves (Gotelli & Colwell 2001) and species-area relationships (Rosenzweig 1995) allow users to assess how different biodiversity indices vary with spatial scale and/or sampling effort. Finally, the package provides functions to simulate sampling processes and to convert the spatially-explicit simulated data into classical community matrices (i.e. sites-by-species abundances matrices). These matrices can then be analysed using standard analytical tools (Legendre & Legendre 2012) to assess how the simulated changes are expressed in measures of biodiversity and influenced by the sampling design. The package is currently available on GitHub (https://github.com/MoBiodiv/mobsim) and also as interactive shiny application with graphical user interface (https://github.com/KatharinaGerstner/mobsimapp). We also plan to publish the package on CRAN (https://cran.r-project.org).

## Package description

### Simulation of community data

An ecological community is characterised by its species-abundance distribution (SAD) and by the spatial distribution of individuals. In `mobsim`, users can use a predefined SAD and add simulated positions of individuals, or simulate both the SAD and the positions (Table 1). For the simulation of SADs, a wrapper around the function `rsad` from the R package `sads` is provided, which offers many options for the underlying statistical distribution (Prado, Miranda & Chalom 2016). In contrast to `sads::rsad`, the function `mobsim::sim_sad` allows the simultaneous specification of the simulated number of individuals and the number of species in the pool. Due to random sampling, there can be fewer species in the simulated community than in the user-defined species pool, but we also provide an argument in `sim_sad` to allow the number of simulated species equal that of the pool size. In this case there can be deviations from the underlying statistical distribution, because rare species are added until the required species richness is reached, leading to a longer tail than expected from the underlying statistical SAD model.

The spatial coordinates of individuals are simulated as 2-dimensional point processes in `mobsim` following Wiegand & Moloney (2014) either using a Poisson process, where individuals are placed randomly, or a Thomas process, where individuals of the same species are clustered. For the Thomas process, users define the numbers and sizes of the clusters, and the number of individuals per cluster, either independently or jointly for all species. The Thomas-process only considers intraspecific aggregation. Individuals of different species are distributed randomly with respect to each other (McGill 2010).

### Analysis of community data

`mobsim` offers several functions to derive spatial and non-spatial patterns from simulated or empirical spatial data. Conversion of empirical data to the format required by `mobsim` is facilitated by auxiliary functions. The function `spec_sample_curve` derives the expected number of species given a certain number of sampled individuals. Individuals are sampled either randomly, giving the well-known species rarefaction-curve (Gotelli & Colwell 2001), or sampling proceeds always from a focal individual to the nearest neighbour, which results in the spatial species-accumulation curve (spatial SAC) (Chiarucci *et al*. 2009). Note that this is different from the sample-based accumulation curve described in Gotelli & Colwell (2001), which considers the distribution of individuals among plots, but not the spatial location of plots.

The function `divar` (diversity-area relationships) estimates several diversity indices for sampling plots of user-defined areas. Accordingly, this function can be used to derive the species-area and endemics-area relationships (Rosenzweig 1995; Harte & Kinzig 1997). In addition, `divar` estimates the Shannon- and Simpson-diversity indices, and their corresponding effective numbers of species (ENS) (Jost 2006).

The function `dist_decay` estimates the distance decay of community similarity (Nekola & White 1999; Morlon *et al*. 2008). The calculation is based on pairwise similarity indices of species composition in square sampling plots using the function `vegdist` from the R package `vegan` (Oksanen *et al*. 2017). `mobsim` includes standard plotting functions for community data and for all the biodiversity patterns introduced. See the online documentation for detailed information on all main and auxiliary functions.

### Sampling of community data

`mobsim` also provides functionality to simulate sampling processes by distributing square plots in a community. The data type provided by the sampling is a sites-by-species matrix, which is a classical data type in community ecology (Legendre & Legendre 2012). Users can choose the size and number of sampling quadrats, as well as the spatial design.

**Table 1:** List of main functions in `mobsim`

| Function name | Category | Description |
| --- | --- | --- |
| `sim_sad` | simulation | Simulate species-abundance distributions (SADs) |
| `sim_poisson_coords` | simulation | Add spatially random coordinates to a SAD |
| `sim_thomas_coords` | simulation | Add coordinates with intraspecific aggregation to an SAD |
| `sim_poisson_community` | simulation | Simulate a community with certain SAD and spatially random coordinates |
| `sim_thomas_community` | simulation | Simulate a community with certain SAD and intraspecific aggregation |
| `spec_sample_curve` | analysis | Derive rarefaction and individual-based species-accumulation curves |
| `divar` | analysis | Derive diversity-area relationships from square sampling plots of different sizes. Diversity indices include species richness, no. of endemics, Shannon index, Simpson index and the effective numbers of species based on Shannon and Simpson indices |
| `dist_decay` | analysis | Derive the distance-decay function from pairwise community similarity measures of square sampling plots |
| `sample_quadrats` | sampling | Virtual sampling of different communities using square plots of user-defined sizes. Users can choose different sampling designs, including random sampling, transect and lattice designs. |

**Figure 1:**
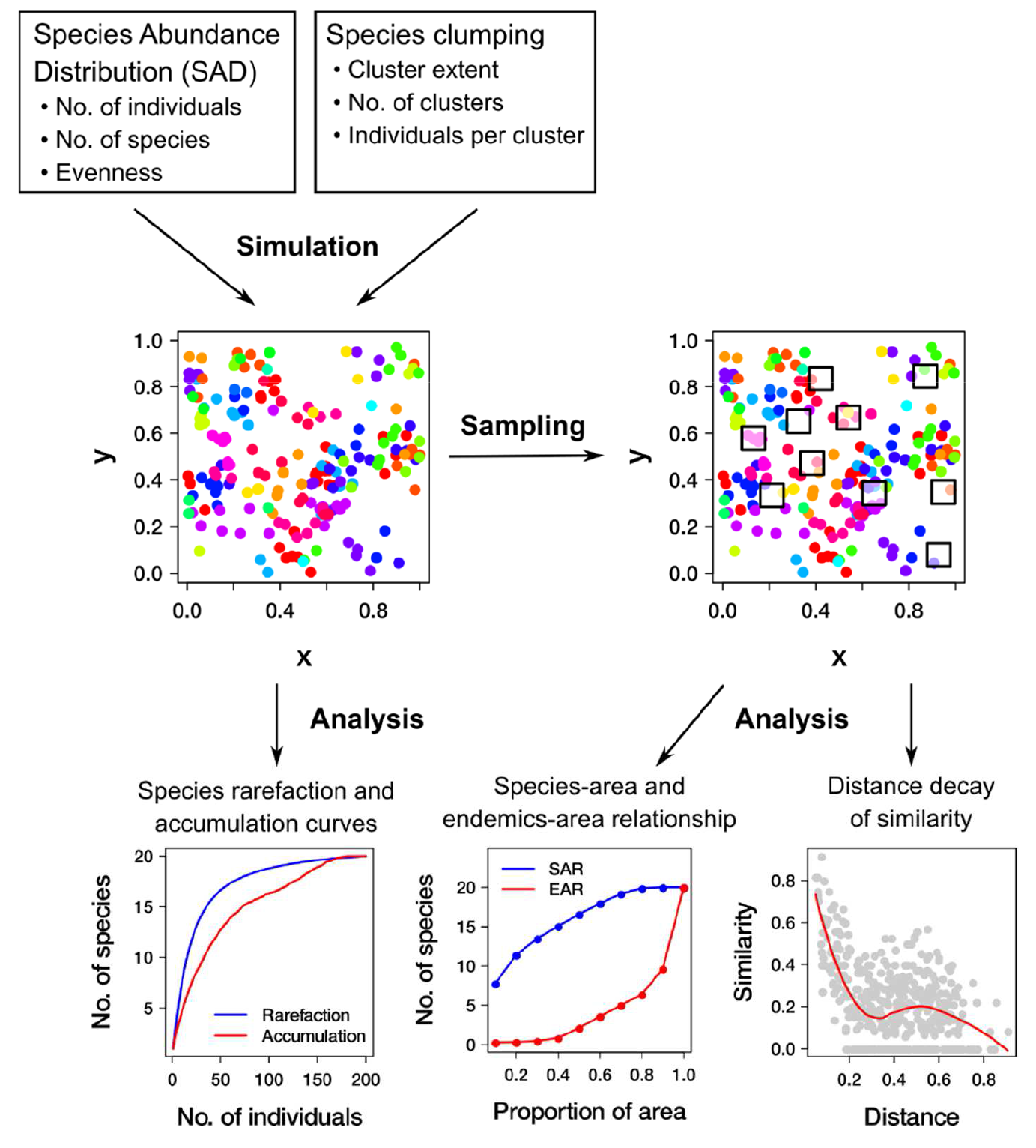
General overview on the purposes of `mobsim`. The package provides functions to simulate species abundances and spatial distributions based on user-defined parameters, including the numbers of species and individuals, the species-abundance distributions, and the aggregation of conspecific individuals. Simulated or observed distributions can be analysed and visualised using `mobsim` functions. Alternatively, sampling processes can be simulated using `mobsim` and the analysis can be done with additional software for classical site-by-species abundance matrices.

## Example applications

Here we present two example applications of `mobsim`: (i) on changes of biodiversity components and (ii) an assessment of species richness estimators. A third example, on extinctions due to habitat loss is provided in the online supporting information.

### Changes of single biodiversity components

The biodiversity in a sampled area depends on three components that can vary independently: (1) the total number of individuals, (2) the SAD of the species pool, and (3) the spatial distribution of individuals and species (McGill 2011; Chase & Knight 2013). Using simulations, we show how the combination of rarefaction and accumulation curves can be used to disentangle changes in these three biodiversity components (Fig. 2). The key point is that the shape of the rarefaction curve only depends on the underlying SAD, but not on the spatial distribution (Gotelli & Colwell 2001), while the shape of the accumulation curve depends on both the SAD and the spatial distribution. First, we randomly removed 50% of all individuals. This does not affect the underlying SAD, as indicated by overlapping rarefaction and accumulation curves that end at different numbers of individuals (Fig. 2a, b). Second, we simulated communities with lower evenness by increasing the variation of species abundances. This resulted in changes in both the rarefaction and the accumulation curves (Fig. 2c, d). Here, the difference (or ratios) between the curves changes with sampling effort, which indicates scale and/or sampling effort dependent effect sizes (Chase & Knight 2013). Despite having the same number of species in the pool, the simulated communities differ in species richness even for the maximum number of individuals, because the rarest species are not sampled into the local community. See Fig. S1 for the same figure that used a fixed species richness of the simulated community, where the curves converge at the largest number of individuals. Third, we add intraspecific aggregation by using a Thomas-process instead of a Poisson-process, which does not affect the rarefaction curve, but leads to lower expected species richness in the spatial species-accumulation curve (Fig. 2e, f).

**Figure 2:**
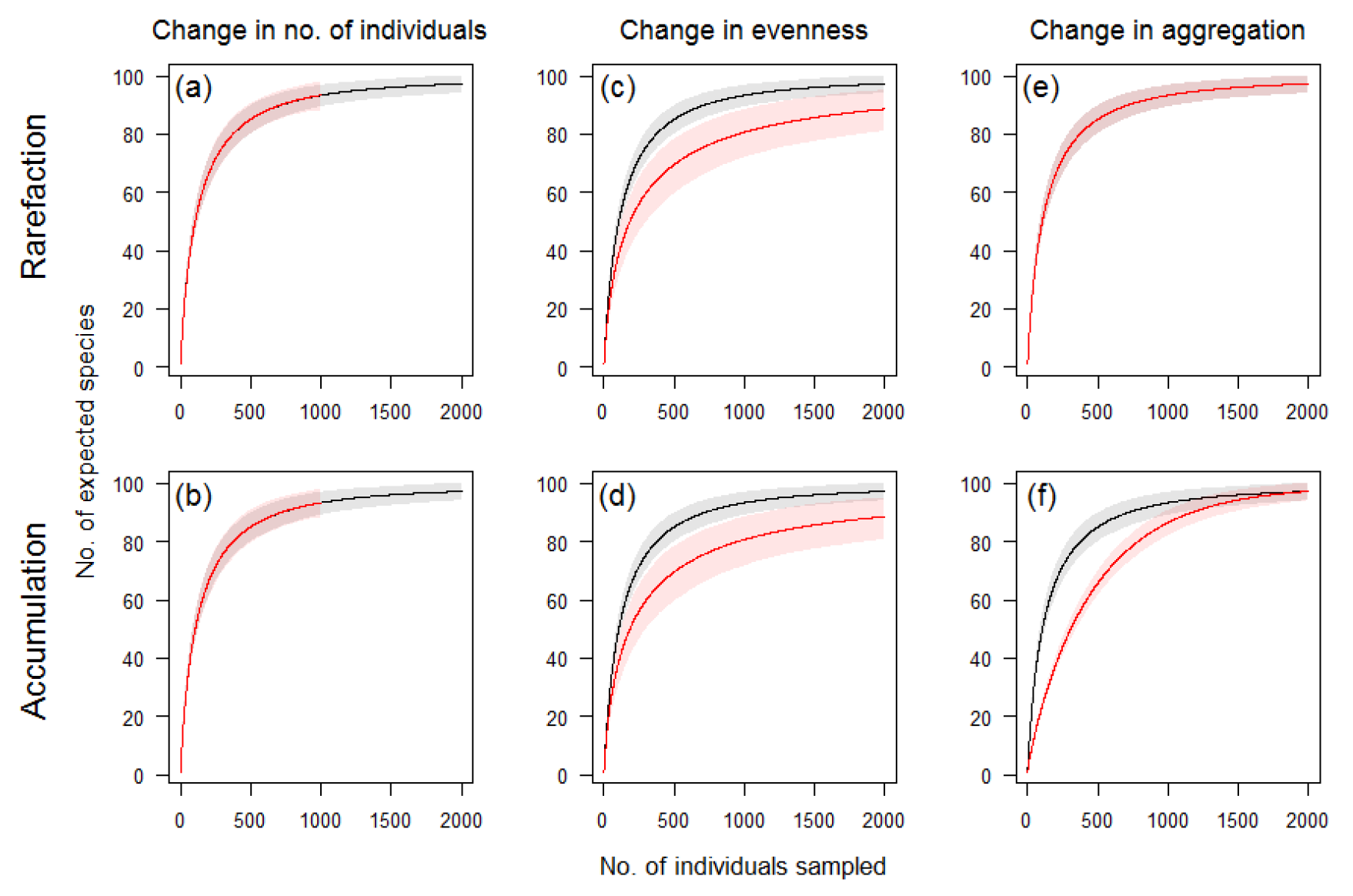
Simulated species rarefaction- and accumulation-curves for changes of three different biodiversity components. The black lines and intervals show the reference community with 2,000 individuals and 100 species in the species pool, a log-normal species-abundance distribution (SAD) with `meanlog = 3` and `sdlog = 1`, and spatially random positions (Poisson distribution). The red lines show changed communities for half the number of individuals (first column), a decrease in evenness (second column), simulated as higher variation in abundances with `sdlog = 1.5`, and a higher intraspecific aggregation (third column), simulated with a Thomas-process with cluster extent `sigma = 0.02`. Ribbons indicate 95% confidence intervals derived from 1,000 replicate simulations.

### Testing species-richness estimators

We are often interested in inferring biodiversity and/or its change in a larger region based on a limited amount of samples. Species-richness estimators offer approaches for estimating the biodiversity of a large community based on local samples (Colwell & Coddington 1994; Chiu *et al*. 2014). However, it remains an open question of how well these estimators perform for different communities and for different sampling strategies (Colwell & Coddington 1994; ter Steege *et al*. 2017). The simulation tools of mobsim are well-suited to address this issue.

We used `mobsim` to assess the performance of a bias-corrected version of the well-known Chao1-estimator (Chiu *et al*. 2014), in the face of spatial aggregation and different sampling designs. We simulated a community with 1,000 species and 1,000,000 individuals. Then we used the function `sample_quadrats` to sample from the community. We varied the proportion of total area sampled between 0.01% - 1% as well as the number of sampling quadrats (1 − 100) that jointly represent the total sampling effort. These combinations of sampling strategies were applied to communities with the same SAD, but with different intraspecific aggregations. We examined four scenarios for aggregation: (1) a random distribution; (2) several large clusters per species; (3) several small clusters per species; (4) one large cluster per species. We used the function `vegan: estimateR` to calculate the species-richness estimator of Chiu *et al.* (2014).

For the community with a random distribution we found no influence of whether a single large or several small quadrats were sampled (Fig. 3a). However, the estimated richness and its uncertainty strongly varied with total sampling effort. The bias of the estimator decreased with increasing sampling effort, but the estimated and true values only converged at the highest effort. At the same time, the uncertainty decreased drastically with sampling effort (Fig. 3a).

For aggregated distributions, the spatial configuration of sampling mattered and a sampling strategy with several small quadrats was less biased than few large quadrats (Fig. 3b-d). For high aggregation, species richness and its uncertainty were strongly underestimated (Fig. 3c, d). This is an important finding, because aggregated species distributions tend to be the rule in nature (McGill 2010, 2011).

Our simulation results underline the recommendation by developers of species-richness estimators, that the estimated values should be only interpreted as lower bounds (Chao 1987; Chiu *et al.* 2014). Furthermore, our findings indicate that for aggregated species distributions, both sampling design and sampling effort have a large influence.

**Figure 3:**
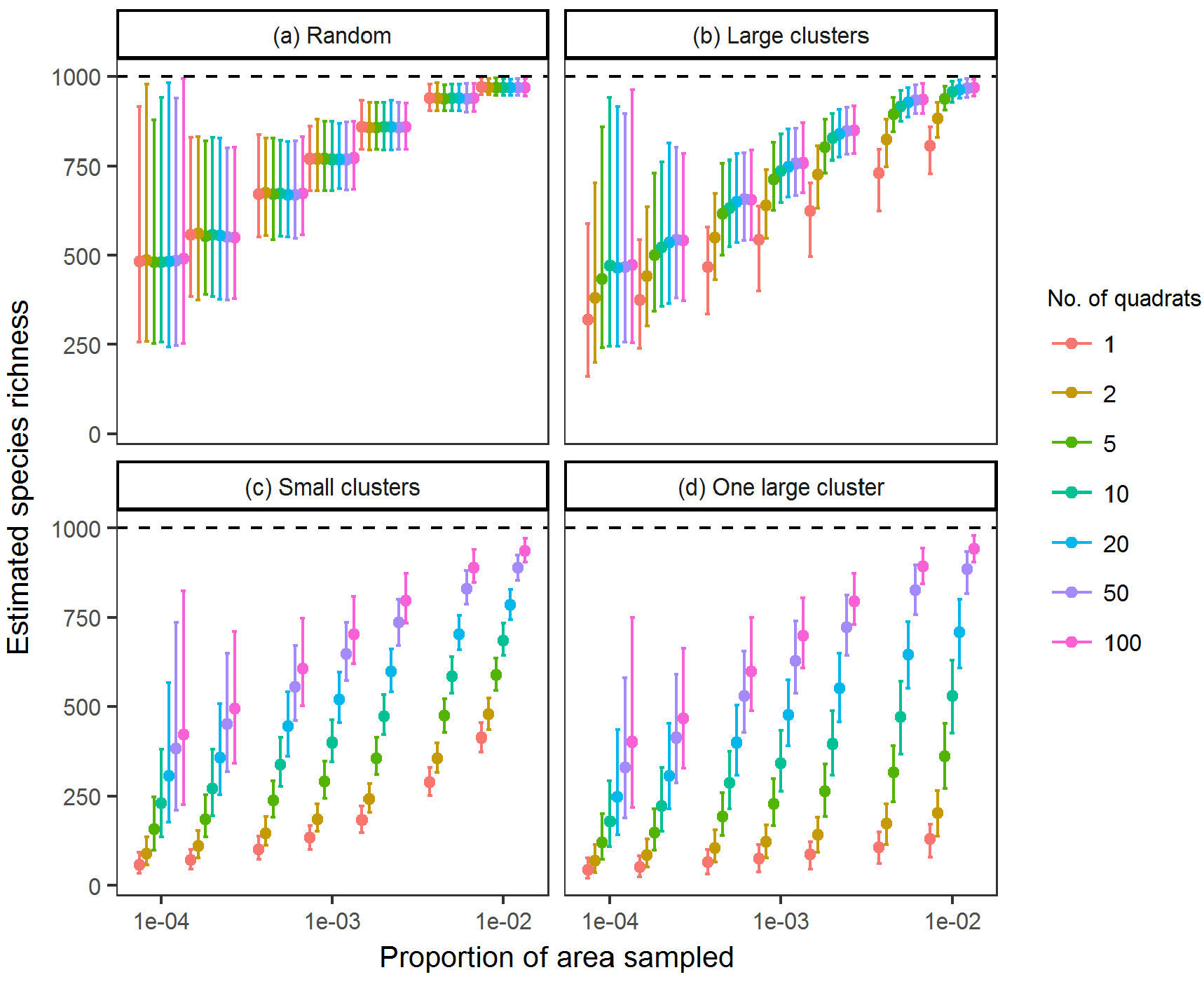
Performance of an asymptotic species-richness estimator for communities with different intraspecific aggregation and for different sampling strategies. The panels show the estimated species richness (chao1 from function `vegan::estimateR`) vs. the proportion of total area sampled. The colour indicate different numbers of randomly distributed sampling quadrats that together form the total amount of sampled area. The different panels show results for communities with the same SAD but different intraspecific aggregation. The points and error bars represent means and 95% confidence intervals from 1,000 replicate simulations. The horizontal lines indicates the true species richness. The following parameter values were used for the simulations: species pool richness: `s_pool = 1000`; number of individuals: `n_sim = 1,000,000`; a log-normal species-abundance distribution (SAD) with `meanlog = 3` and `sdlog = 1`; large cluster size: `sigma = 0.05`, small cluster size: `sigma = 0.01`; one cluster per species: `mother_points = 1`.

## Conclusions

The total number of individuals, the distributions of species relative abundances, and intraspecific aggregation are key components of community structure and any changes in biodiversity are mediated through changes in these components. Furthermore, biodiversity changes are scale-dependent. The combination of tools for simulation and analysis of biodiversity patterns provided in `mobsim` is well-suited to foster understanding on the emergence and consequences of scale-dependent biodiversity changes. The package integrates key tools of community ecology so that ecologists can derive valid and robust interpretations of biodiversity patterns and changes observed in real-world data. We also believe that the use of controlled simulation experiments is highly beneficial for education in biodiversity science.

## Acknowledgements

We gratefully acknowledge the support of the German Centre for Integrative Biodiversity Research (iDiv) Halle-Jena-Leipzig funded by the German Research Foundation (FZT 118).

## Authorship statement

FM and JMC conceived the package concept and structure. FM implemented the first package version. KG, XX, and DM contributed code and supported the package revision. KG implemented the shiny online application. FM wrote the first manuscript draft and all authors critically revised the text and gave final approval for publication.

